# Single cell transcriptome analysis reveals markers of naïve and lineage-primed hematopoietic progenitors derived from human pluripotent stem cells

**DOI:** 10.1101/602565

**Authors:** Antonella Fidanza, Nicola Romanò, Prakash Ramachandran, Sara Tamagno, Martha Lopez-Yrigoyen, Alice Helen Taylor, Jennifer Easterbrook, Beth Henderson, Richard Axton, Neil Cowan Henderson, Alexander Medvinsky, Katrin Ottersbach, Lesley Margaret Forrester LM

## Abstract

During embryogenesis the hematopoietic system develops through distinct waves that generate progenitors with increasing lineage potential, ultimately producing haematopoietic stem cells (HSCs). In vitro differentiation of human pluripotent stem cells (hPSCs) follows the early steps of haematopoietic development but the production of HSCs has proven more challenging. To study the dynamics and heterogeneity of hematopoietic progenitor cells generated in vitro from hPSCs, we performed RNA sequencing of over 10000 CD235a^-^CD43^+^single cells. We identified the transcriptome of naïve progenitors and those primed toward erythroid, megakaryocyte and leukocyte lineages, and revealed their markers by clustering, trajectory analyses and functional assays. CD44 marks naïve clonogenic progenitors that express the transcription factor, LMO4 and can be expanded upon BMP4 stimulation. Naïve progenitors give rise to primed CD326^+^erythroid, ICAM2^+^CD9^+^megakaryocyte, and monocyte, neutrophil and eosinophil progenitors. We have generated an online dataset of human hematopoietic progenitors and their transcriptional remodelling upon lineage priming.

## Introduction

Human pluripotent stem cells (hPSC) can be differentiated in vitro into various cell types, providing both a model for basic research studies and a source of clinically relevant cells (Vo and Daley, 2015). During development, two waves of restricted hematopoietic progenitors arise in the extraembryonic tissues of the yolk sac, before hematopoietic stem cells (HSCs) emerge in the embryo proper (Palis, 2016). The first “primitive” wave, gives rise to erythrocytes, megakaryocytes and macrophages from embryonic day E7.25 in the mouse embryo (Palis *et al.*, 1999; Tober *et al.*, 2007). From E8.25, the first “definitive” progenitors, called erythro-myeloid progenitors (EMPs) constitute the second wave and these can be distinguished from the primitive progenitors by their potential to generate granulocytes (McGrath *et al.*, 2015). Intraembryonic haematopoiesis is established during E10.5-E11.5 by the emergence of HSCs, the only cells that can sustain the lifespan production of all blood lineages, and maintain this property upon transplantation (Medvinsky and Dzierzak, 1996).

Despite many laboratories successfully recapitulating the development of multilineage hematopoietic progenitors from hPSCs in vitro, the robust derivation of bona fide long-term repopulating hematopoietic stem cells (HSCs) has not been achieved (Ditadi, Sturgeon and Keller, 2017). Two main strategies have been employed in attempts to overcome this challenge: modification of culture conditions to mimic embryonic ontogeny and the overexpression of transcription factors in genetic programming experiments (Ivanovs *et al.*, 2017). Because the ontogeny of the human hematopoietic system is poorly characterised, reproduction of the natural molecular cues occurring in the embryo is arduous.

Furthermore, the broad overexpression of target genes identified by bulk approaches has failed to precisely reproduce the transcriptome of HSCs. In addition, the dynamic nature and the heterogeneity of the hematopoietic progenitor cell populations that arise during development poses additional confounders to the identification of both signalling and target genes. To overcome these limitations, we propose that the in-depth characterisation of hPSC-derived hematopoietic progenitors at the single cell level, and the subsequent comparison with data sets obtained from haematopoietic progenitors generated in vivo, will be instrumental for the development of new and improved strategies for their in vitro production. Some single cell expression profiles of hPSC derived hematopoietic cells have been reported to date, but they either used biased approaches such as preselected probes (Guibentif *et al.*, 2017), or used a limited number of cells sorted with multiple markers, thus impacting the heterogeneity resolution and the detection of subpopulations (Angelos *et al.*, 2018).

Combining the use of two reporter cell lines and functional assays we designed a minimal membrane marker strategy that allows the isolation of a broad, heterogeneous population of hPSC-derived haematopoietic progenitor cells. We showed that the CD235a^-^CD43^+^cell population contained definitive progenitors marked by the RUNX1C-GFP reporter (Ng *et al.*, 2016) and excluded KLF1-mCherry-expressing committed erythroid cells. To explore the inherent heterogeneity of this population and to decipher the hierarchy of lineage priming we generated a large single cell RNA-sequencing data set of 11420 CD235a^-^CD43^+^cells. Cell surface markers of progenitors and their lineage-primed descendants were identified and validated using in vitro functional assays. Lineage trajectories predicted by pseudotime analyses were validated using a chimeric cell culture system involving a constitutive ZeissGreen reporter iPSC line. This dataset can be compared to in vivo-sourced human HSC allowing the identification of candidate genes and pathways that could be modulated for efficient in vitro HSC production.

## RESULTS

### Minimal marker strategy for the unbiased isolation of hematopoietic progenitors

To resolve the heterogeneity of the definitive haematopoietic progenitor cell population in an unbiased manner, we used functional assays and reporter cell lines to define a minimal marker approach for their isolation from differentiating iPSCs.

CD235a (Glycophorin A), a broad erythroid lineage marker, has been reported to mark in vitro the emerging mesoderm specified to primitive human haematopoietic cells (Sturgeon *et al.*, 2014) and to be retained on primitive erythro-megakaryocytes progenitors (Vodyanik *et al.*, 2006). To study the identity of CD235a expressing cells in our culture system, we sorted CD235a^-^and CD235a^+^cells from differentiating iPSCs at day 13 and analysed their haematopoietic potential in clonogenic assays (Figure 1A, Supplementary Fig 1A). The vast majority of robust colony forming units (CFU-Cs) were generated from the CD235a^-^population, while the CD235a^+^cell population generated largely primitive erythroid colonies and only a few myeloid colonies (Figure 1A), in line with previous reports. Gene expression analyses revealed that CD235a^-^cells expressed higher levels of progenitor-associated genes, including *RUNX1C, TAL1, and GATA2* and lower levels of erythroid markers (*KLF1, GATA1* and *HBE*) compared to CD235a^+^cells (Figure 1B). To verify that the CD235a^-^population consisted of definitive, rather than primitive, haematopoietic progenitor cells we used the RUNX1C-EGFP reporter cell line that marks emerging haematopoietic progenitors (Supplementary Fig 1H-I)(Ng *et al.*, 2016). The *RUNX1* gene encodes different protein isoforms, differentially expressed during development, with RUNX1C being expressed by the distal promoter in definitive cells in vitro and in vivo (Sroczynska *et al.*, 2009; Ng *et al.*, 2016). As predicted, RUNX1C-GFP^+^cells were almost entirely found within the CD235a^-^compartment (Figure 1C) and were marked by CD43 membrane expression (Figure 1D), previously reported to mark human PSC-derived progenitors (Vodyanik *et al.*, 2006; Garcia-Alegria *et al.*, 2018). When isolated by flow cytometry, RUNX1C-GFP^+^demonstrated higher number of CFU-Cs compared to the RUNX1C-GFP^-^population (Figure 1E).

**Figure 1.**
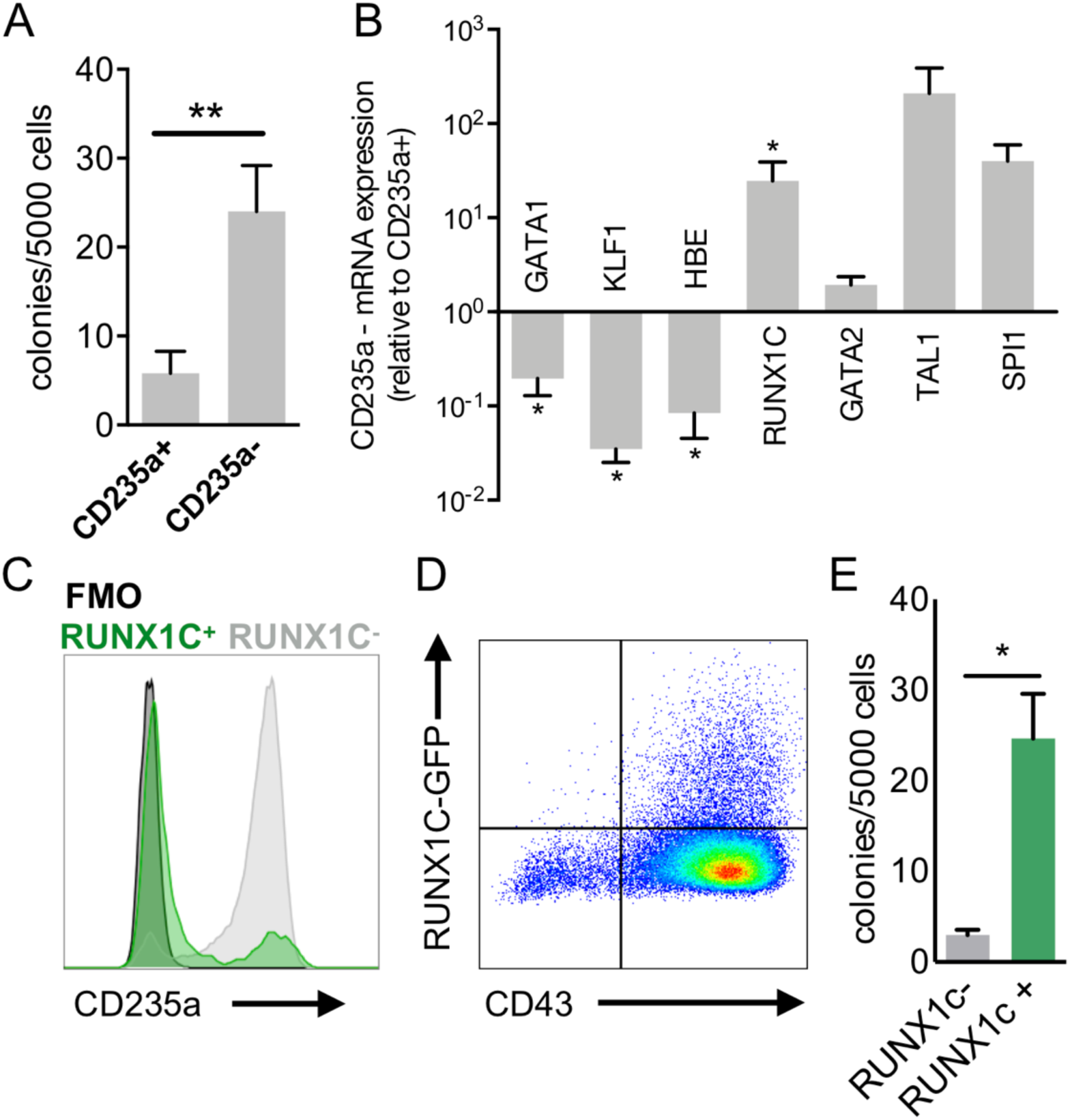
Human definitive hematopoietic progenitors resides in the CD235a^-^CD43^+^compartment. (**A**) Colony forming potential of sorted CD235a+ and CD235a-cells (n=5, p<0.01, paired t-test). (**B**) Gene expression profile of CD235a-cells, relative to CD235a+ (n=6, *p<0.05, Wilcoxon test).(**C**) Distribution of RUNX1C+ and RUNX1C-in relationship to CD235a. (**F**) RUNX1C-GFP expression in relation to CD43. (**G**) Colony forming potential of sorted RUNX1C+ and RUNX1C-cells (n=3, p<0.05, paired t-test).

We confirmed the primitive erythroid bias of the CD235a^+^population by flow cytometry analysis of ε-globin expression, a marker associated with the first wave of erythroid cells (Supplementary Fig 1B). In the murine embryo, definitive erythroid cells enucleate more efficiently than their primitive counterparts, that fully enucleate only after persisting in the bloodstream for several days (Kingsley *et al.*, 2004; McGrath *et al.*, 2008). Thus, to further confirm the primitive bias of CD235a^+^cells, we cultured isolated CD235a^+^and CD235a^-^cells in erythroid differentiation conditions for 17 days and assessed the enucleation efficiency. Consistent with published reports (Olivier *et al.*, 2016; Yang *et al.*, 2017), enucleation efficiency of iPSC derived erythroid cells is very low, but nonetheless we observed that the erythroid cells derived from the CD235a^-^population had a higher potential to enucleate than those derived from CD235a^+^cells (Supplementary Fig 1C). The expression of the transcription factor KLF1, is initiated prior to erythroid commitment and is maintained throughout erythropoiesis (Siatecka and Bieker, 2011), thus, we used a KLF1-mCherry reporter to track erythroid commitment during differentiation (Supplementary Fig 1D). KLF1-mCherry^+^reporter was expressed in cells associated with small primitive erythroid colonies and restricted to the CD235a^+^cells (Supplementary Fig 1E-F). KLF1-mCherry-expressing cells showed a limited colony forming potential compared to the KLF1^-^population (Supplementary Fig 1G).

Taken together these data indicate that CD235a^-^CD43^+^cells, from differentiating hPSCs, contains definitive hematopoietic progenitors and excludes cells derived from the first, primitive wave. We anticipated that CD235a^-^CD43^+^compartment would also comprise the early stages of lineage priming, capturing the hierarchy of early human developing progenitors.

### Single cell RNAseq of iPSCs-derived haematopoietic progenitor cells reveals the transcriptome of naïve and lineage primed progenitors

We collected exclusively suspension cells from two independent replicate cultures at day 13 of differentiation, isolated the CD235a^-^CD43^+^cells containing the definitive haematopoietic progenitors and subjected them to microfluidic single cell RNA libraries preparation followed by sequencing and data analyses (Figure 2A). After quality controls and filtering of the data we obtained the transcriptome of 11420 cells (Supplementary Figure S1A-E). Following dimensionality reduction through Principal Component Analysis (PCA), we used graph-based clustering analysis and obtained 9 clusters of cells (Butler *et al.*, 2018a) and visualised on the tSNE projection (Figure 2B). Although the two replicates did not show obvious differences (Supplementary Figure S2E), we regressed out the batch effect before pulling the samples together for further analysed. We assigned cell identities based on the expression of known markers and identified markers from the dataset that were cluster specific (Figure 2C-D). Clusters containing more immature, unprimed progenitors were identified by their high level of progenitor-associated genes such as *KIT* and *GATA2* and their lack of expression of genes associated with specific cell lineages, and were annotated as naïve populations (Figure 2D). Clusters that displayed expression of lineage markers were annotated as primed progenitors (Figure 2B-D). These included clusters of cells primed towards the megakaryocyte (*GP9* and *PF4*), erythroid (*GYPA* and *KLF1*) and granulocyte (*AZU1* and *PRNT3*) lineages (Figure 2D). Markers for each of the cell clusters were identified by differentially expressed gene analysis and further supported the identities assigned to of these clusters (Figure 2C and Supplementary Table S1).

**Figure 2.**
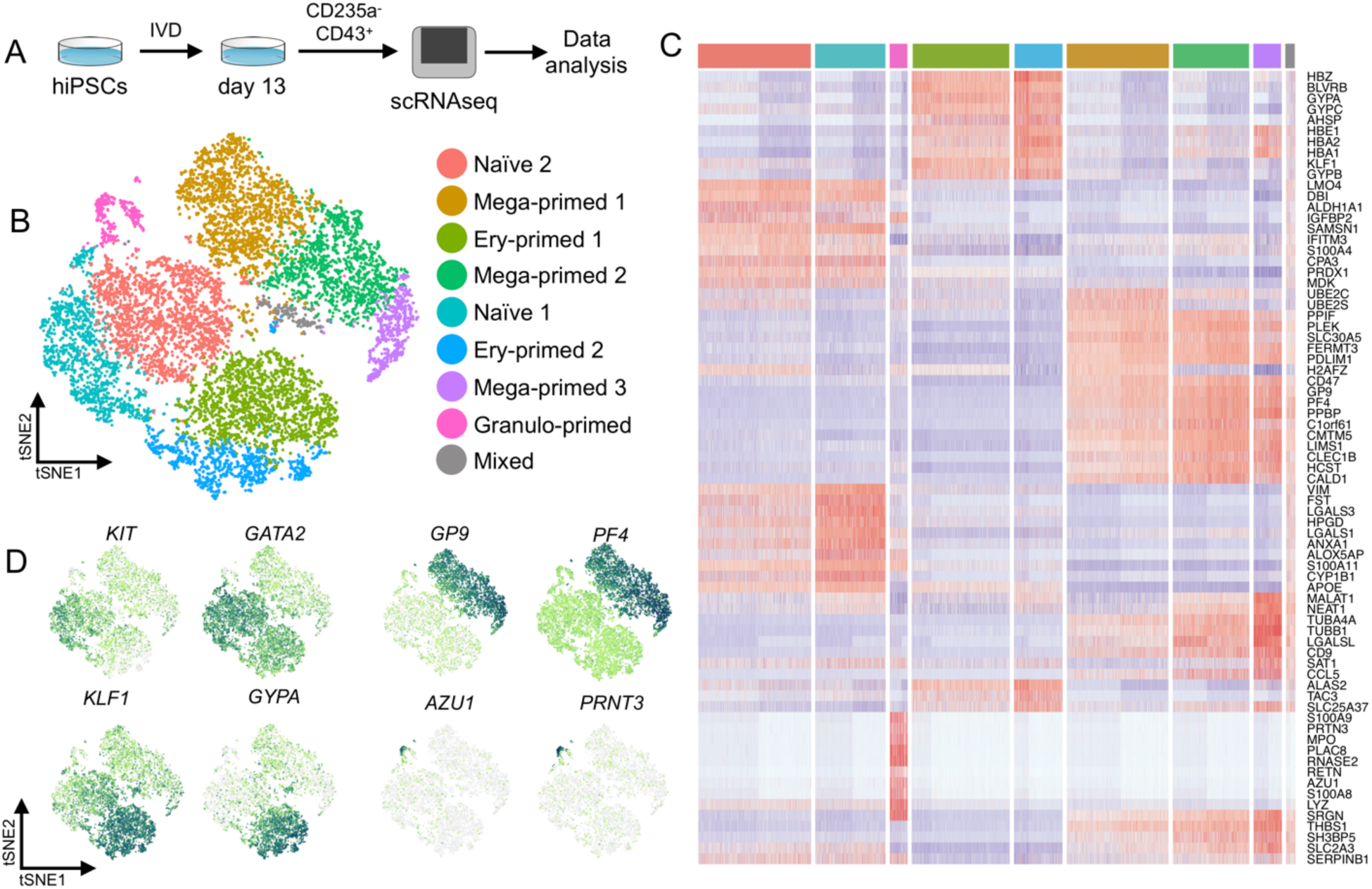
Single cell transcriptome analysis identifies cluster of progenitors and primed cells. (**A**) Schematic of the single cell RNA sequencing experiment. (**B**) tSNE visualization of 11,420 cells divided into 9 clusters. (**C**) Heatmap of the top 10 marker genes for each cluster. (**D**) Gene expression of cell type markers on tSNE.

### Trajectory analyses reveal the hierarchy of in vitro derived hematopoietic progenitors

To study the hierarchical relationship between cell populations we performed trajectory analysis using two different methods: diffusion analysis (Haghverdi *et al.*, 2016) using the Seurat R package (Butler *et al.*, 2018b) and pseudotemporal ordering, using the Monocle R package (Qiu *et al.*, 2017) (Figure 3A-D). Diffusion analysis identified a central core from which three distinct trajectories appeared to emerge. The central core corresponded to cells belonging to the progenitor clusters that we had annotated as naïve 1 and naïve 2 (Figure 3A-B). Branches comprised cells that expressed genes associated with specific lineages, that we annotated as Ery-, Mega-and Granulo-priming directions. These three lineages are expected to branch from EMPs, progenitors of the second wave of yolk sac haematopoiesis (McGrath *et al.*, 2015). Comparable trajectories were observed using pseudotemporal ordering. After calculating a pseudotime value for each cell, we ordered them starting from a root state corresponding to the branch containing cells that were located at the core of the diffusion plot and that we had identified as naïve progenitors in our original clustering (Figure 3C). Pseudo time reconstruction of the hierarchy showed that cells we had annotated as naïve 1 were located at the top of the hierarchy and appeared to progress to naïve 2 cells before entering branches that consisted of lineage primed cells (Figure 3C-D). Lineage priming was also inferred by the expression of lineage-associated transcription factors (Figure 3E) that were identified by filtering of marker genes according to their GO annotation. For instance, erythroid primed clusters demonstrated expression of both *KLF1* and *MYC* (Figure 3E), with the expression level of the latter decreasing in Ery 2 compared to Ery1, according to their position within the differentiation hierarchy (Figure 3D). For mega-primed cluster 1 and 2 we observed the expression of *GATA1, TAL1* and *FLI1* (Figure 3E), a cocktail of genes recently used for forward programming of hiPSCs to the megakaryocyte lineage (Moreau *et al.*, 2016). Granulo-primed cells were represented by a separate branch and showed the expression of *CEBP-D, CEBP-B, CEBP-A* and *CEBP-E* (Figure 3E).

**Figure 3.**
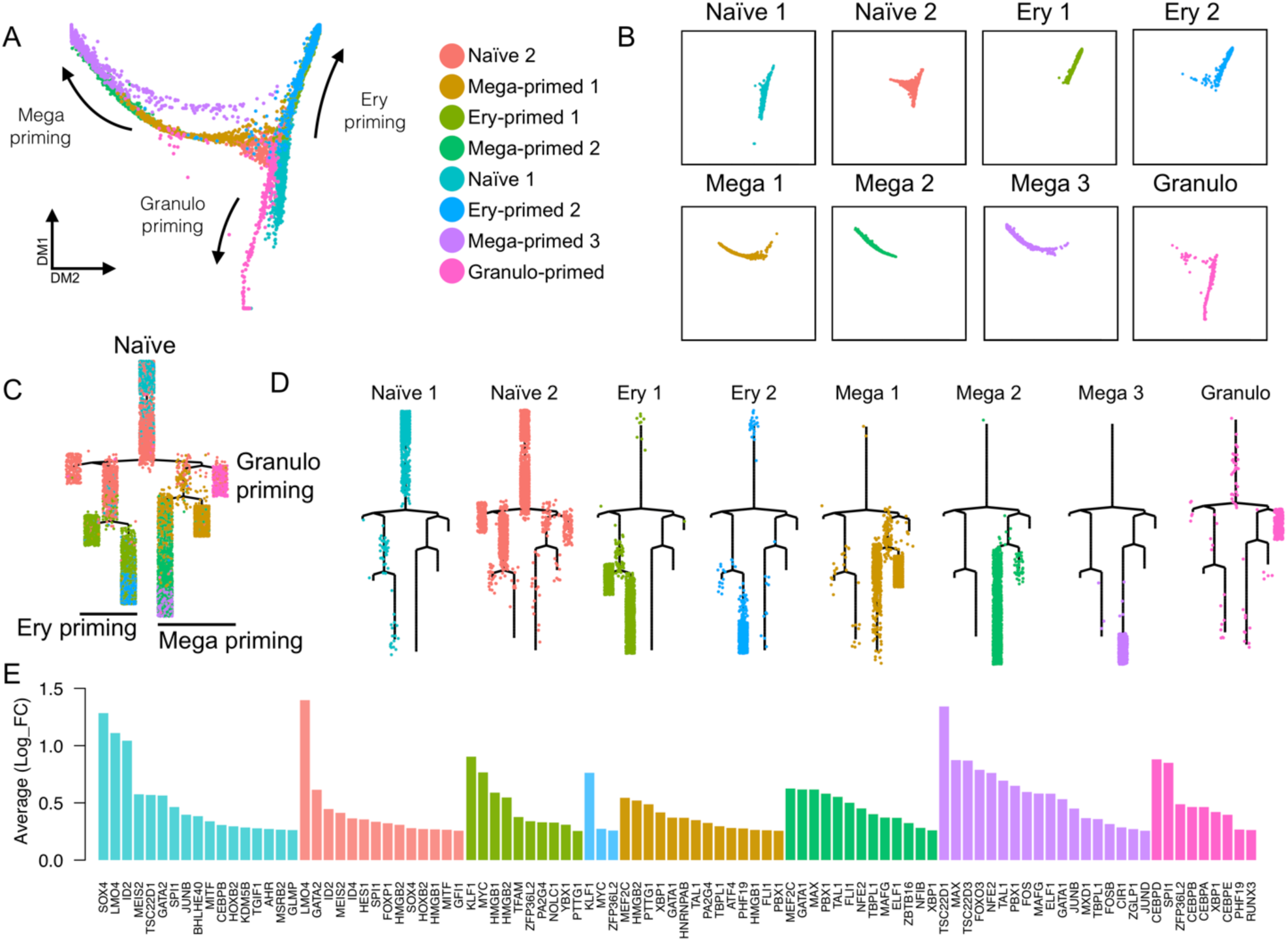
Trajectory analyses confirm progenitors identity and outlines priming directions. (**A**) Diffusion analysis plot shows progenitor in the core region with primed cells describing the different direction of priming. (**B**) Representation of single clusters on the diffusion plot. (**C**) Monocle trajectory analysis with shows same priming obtained from the diffusion plot, (**D**) single cluster visualized on trajectory. (**E**) Top transcription factors expression in each cluster.

### CD44 membrane expression marks clonogenic human hematopoietic progenitors

To functionally validate the results of our trajectory analyses, we assessed the haematopoietic potential of cells that we had defined as naïve progenitor populations. We filtered the list of marker genes encoding membrane proteins and designed a prospective sorting strategy to isolate the progenitor populations. Genes encoding the cell surface markers *CD33, CD44*, and *ITGB2* (also known as CD18) were enriched within clusters associated with the naïve progenitors and so we hypothesised that these markers could be used for their isolation (Supplementary Figure 3). CD33 was expressed uniformly by both naïve 1 and naïve 2 populations whereas CD44 and CD18 expression appeared higher in the naïve 1 population (Supplementary Figure 3). We used CD44 and CD18 to fractionate the CD235a^-^CD43^+^CD33^+^cell population and identified the naïve 1A (CD44^+^CD18^-^), naïve 1B (CD44^+^CD18^+^) and naïve 2B (CD44^-^CD18^-^) populations (Figure 4A). Trajectory analysis predicted that the cell population defined as naïve 1 was at the top of the hierarchy and gave rise to the naïve 2 cell population prior to lineage priming (Figure 3C-D). To functionally test this prediction, we used a chimeric co-culture system where input cells can be tracked within differentiation conditions (Figure 4B). We synchronously differentiated the SFCi55-ZsGreen iPSC line, that expresses the Zeiss Green reporter in a constitutive manner (Lopez-Yrigoyen *et al.*, 2018), and their parental SFCi55 line. To verify the progression from naïve 1 to naïve 2 cells and from naïve 2 to lineage primed cells detected at day 13, we sorted the naïve 1 cells (CD33^+^CD44^-^CD18^-^) and naïve 2 (CD33^+^CD44^+^CD18^-/+^) from the SFCi55-ZsGreen iPSCs at day 10 and then co-cultured them with the synchronised parental SCFi55 differentiating iPSCs cells up to day 13. As predicted from the trajectory analysis, naïve 1 cells were able to generate ZsGreen-expressing naïve 2 cells in the chimeric culture. We also noted that naïve 1 cells also retained their immunophenotype, indicating some self-renewal capacity (Figure 4C). Interestingly, naïve 2 cells demonstrated some potential to acquire the naïve 1 markers, CD44 and CD18 (Figure4C), suggesting a degree of fluidity between the progenitors’ compartments. As predicted by the trajectory analysis (Figure 3 A-D), ZsGreen-expressing naïve 2 cells acquired also the ability to generate haematopoietic cell progeny as predicted by the trajectory analysis (Figure 3 A-D) comprising erythroid cells (CD235a^+^), megakaryocytes (CD41^+^) and adherent mature macrophages (25F9+) (Supplementary Figure 3B). We compared the CFU-C capacity of naïve 1 and 2 progenitors that were present at the different stages of the differentiation protocol. When plated in clonogenic methylcellulose assays, only cells expressing CD44 on their membranes, naïve 1 cells isolated from day 10 and 13, formed CFU-C colonies, whereas virtually no colonies were generated by naïve 2 cells (Figure 4D-E). These data indicate that CD44 membrane expression alone resolves CFU-C forming cells and supports the hierarchy predicted by the trajectory analyses (Figure 3A-D). Our results demonstrate that our chimeric co-culture system is able to assess the lineage output that cannot be assessed by methylcellulose assays alone. To assess whether the naïve cell populations identified using our unique sorting strategy showed features of definitive haematopoietic progenitors, we assessed the expression of the RUNX1C-GFP reporter in these cells. We observed RUNX1C-GFP expression in both cell types, with a higher proportion of RUNX1C^+^cells in the naïve 1 compared to naïve 2 population (Supplementary Figure 3C). We then focused our attention on the other transcription factors expressed by the progenitor clusters and identified high levels of *ID2* and *ID4* in naïve progenitors (Figure 3E), with *ID2* highly expressed in naïve 1. ID genes are targets of BMP signalling, so we predicted that naïve populations could be modulated by addition of BMPs. To test this hypothesis, we included BMP4 in the differentiation culture from day 10, when both naïve 1 and 2 were present and then assessed the proportion of these cells by day 13. The frequency of both naïve 1 and 2 cells increased by 25% and 59% respectively in the presence of BMP4 indicating that this signalling pathway was involved in their expansion (Figure 4F).

**Figure 4.**
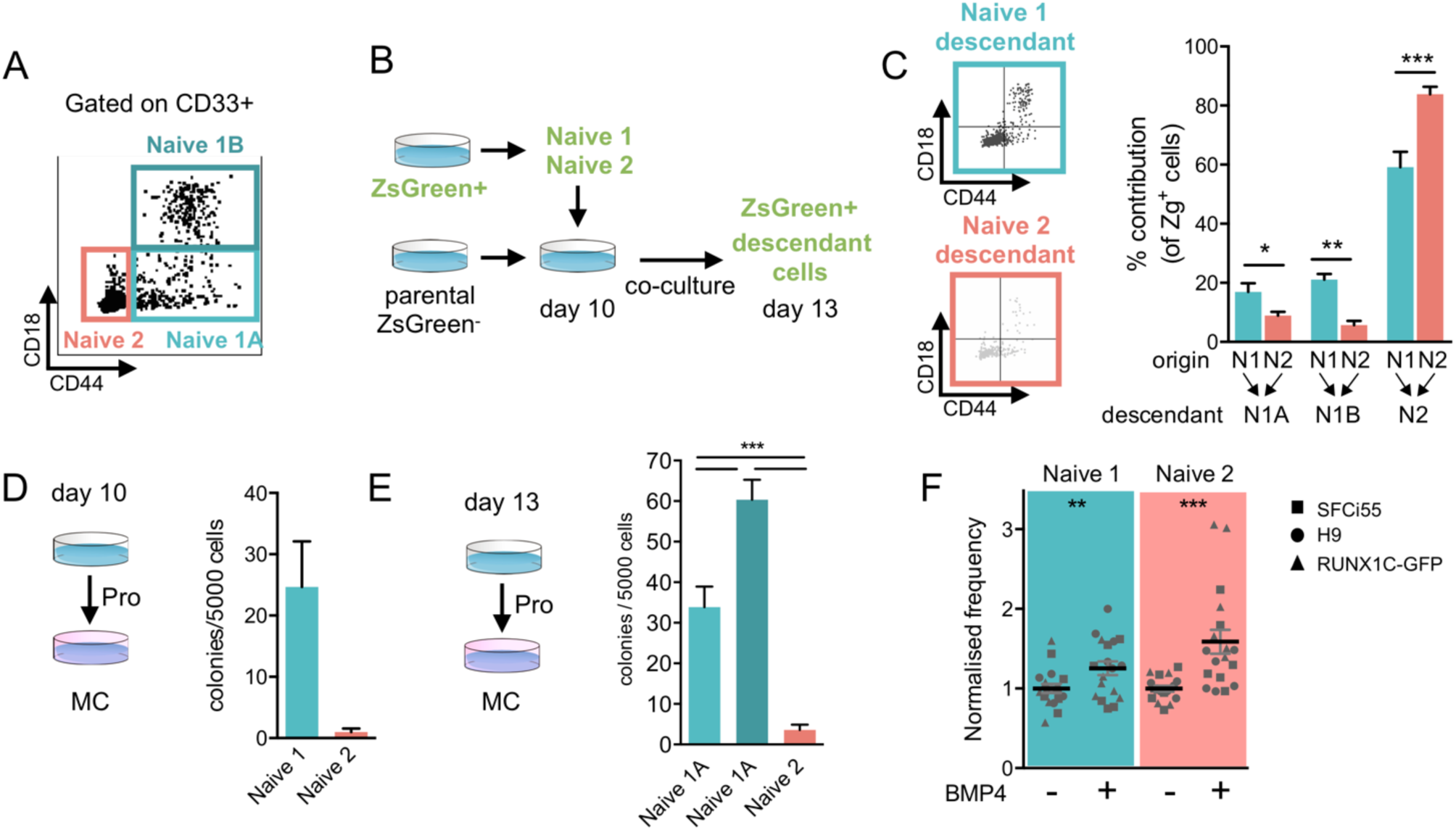
CD44 identifies clonogenic hematopoietic progenitors that can be expanded upon BMP4 addition. (**A**) Scatter plot of flow cytometry profile of Naïve 1, 2A and 2B cells are gated on CD235a^-^ CD43^+^CD33^+^. (**B**) Schematic of the chimeric culture system to trace cells during the differentiation. (**C**) Contribution of Naïve 1, in teal, and Naïve 2, in pink, to the Naïve 1, 2A and 2B compartment (n=6, multinomial logistic regression, *p<0.05, **p<0.01, ***p<0.005). (**D**) Colony forming assay result for Naïve 1 and Naïve 2 cells sorted at day 10 (n=3, paired t-Test p=0.0753) *CD44* expression distribution in Naïve 1 and in Naïve 2 cluster. (**E**) Colony forming assay result for Naïve 1, 2A and 2B cells sorted at day 13 (n=9, Holm-Sidak’s test, p<=0.001). (**F**) Frequency of Naïve 1 and Naïve 2 cells upon addition of BMP4 (n=6 for each cell line, mixed effects model, **p<0.005,***p<0.001).

### CD44 and LMO4 are co-expressed in definitive hematopoietic sites in the mouse embryo

We identified CD44 as a marker of naïve progenitors with tri-lineage potential ascribable to the EMPs progenitors that, in the mouse model, reside in the yolk vasculature. To test if this marker was labelling hematopoietic clusters in the mouse yolk sac we assessed CD44 expression by immunostaining and flow cytometry (Figure 5A-B). At E10.5, CD44 was expressed on endothelial cells in a bimodal pattern, with vessels expressing low and high levels, with the latter containing very bright clusters of haematopoietic cells (Figure 5B). By flow cytometry, we observed that by E11, CD44^high^cells contained all the CD45^+^cells and a proportion of VeCad^+^(Figure 5A). Within the embryo proper, CD44 was expressed on the membrane of endothelial cells within the dorsal aorta, contrary to the venous endothelial layers that were negative for CD44 (Figure 5C). By flow cytometry we observed that CD44 was co-expressed with CD45^+^in the AGM region. These data suggest that CD44 is expressed on haemogenic endothelial cells and it is retained on the hematopoietic cells arising from it.

**Figure 5.**
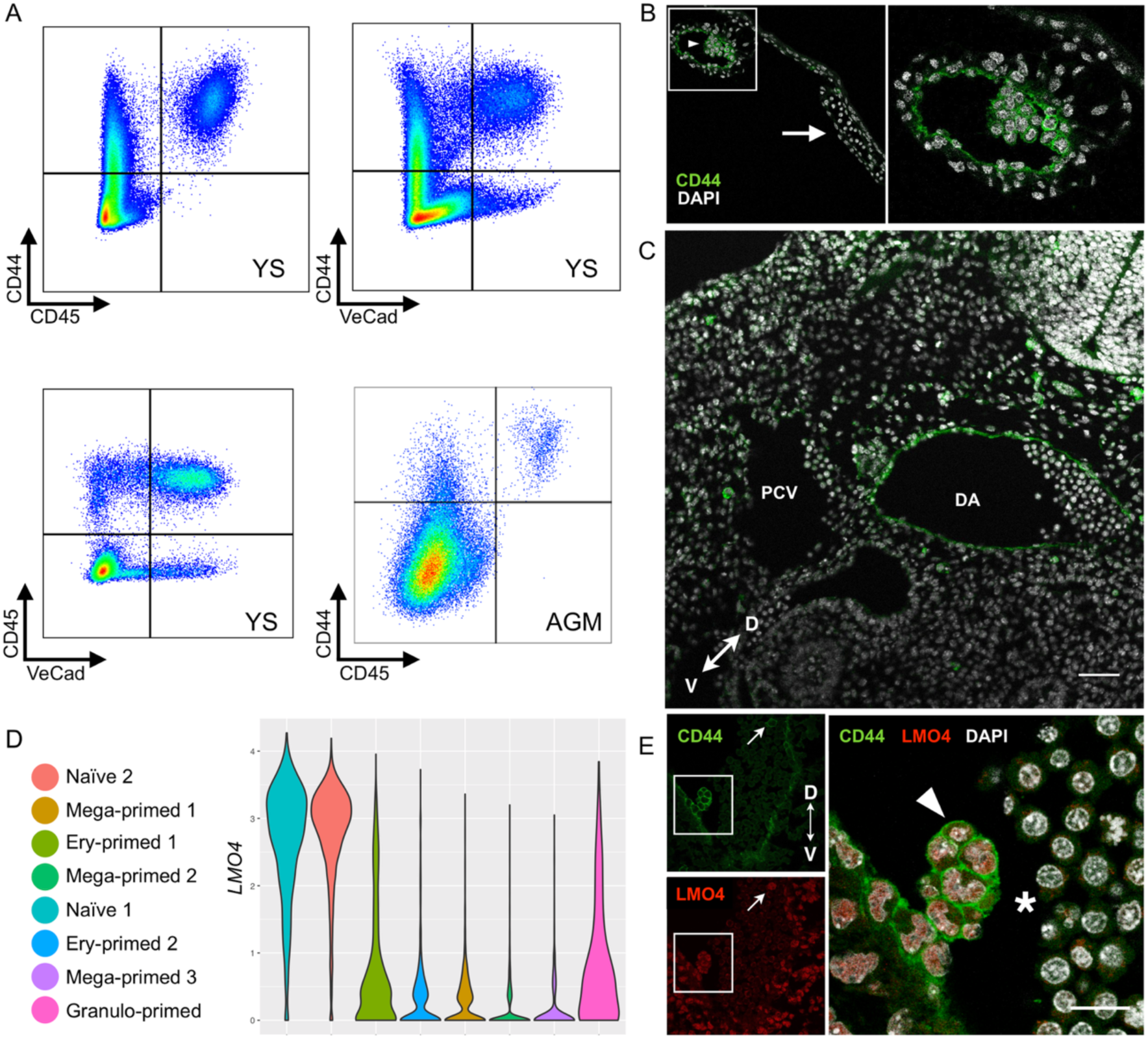
CD44 and LMO4 is co-expressed in human and mouse definitive progenitors. (**A**) Flow cytometry of E11 yolk sac showing expression of CD44 in relation to CD45 and VeCad, and AGM region at E10.5. (**B**) CD44 immunostaining in E10.5 yolk sac (arrowhead: CD44+ vessel and hematopoietic cluster, arrow: CD44-vessel). (**C**) CD44 immunostaining of the dorsal region of the mouse embryo at E10.5 showing expression on arterial endothelial cells (DA - dorsal aorta, PCV - posterior cardinal vein, bar=50μm). (**D**) LMO4 gene expression distribution across clusters. (**E**) LMO4 and CD44 immunostaining in the AGM region (transverse section, zoomed image on intra-aortic cluster bar=15μm, arrow: circulating cell CD44+LMO4+, arrowhead: IAHC, asterisk: circulating negative cells).

From our single cell RNA profiles, we observed that the transcription factor *LMO4* was expressed in cells within the progenitor clusters and subsequently downregulated upon lineage priming (Figure 3E and 5D). As this transcription factor has not been associated previously with definitive haematopoietic progenitors, we assessed its expression during haematopoietic development in vivo. We immunostained sections of the aorta-gonad-mesonephros (AGM) region of the developing E10.5 mouse embryo where intra-aortic hematopoietic clusters (IAHC) are mainly composed of hematopoietic progenitors and pre-HSCs (Figure 5E, supplementary Figure 3D). CD44 was highly expressed by cells of the IAHCs and by single circulating cells (Figure 5E, supplementary Figure 3D) and at lower levels by the aortic endothelium. LMO4 was co-expressed at high levels in the nuclei of cells with CD44 on their membrane, both in IAHC cells and in rare circulating cells (Figure 5E, supplementary Figure 3D). Neither CD44 nor LMO4 were expressed in the majority of circulating cells within the lumen of embryonic vessels (Figure 5E), which at this stage of development are mainly primitive erythroid cells.

### Identification of membrane markers of lineage primed progenitors

We identified clusters with lineage primed signatures and selected membrane markers that we predicted could be used for their isolation. Although our original sorting strategy excluded CD235a^+^erythroid cells, we nevertheless detected erythroid primed progenitors (Figure 3B). Ery-primed clusters 1 and 2 both showed expression of *EPCAM* and *MYC* (Figure 3E, 6A), indicative of early committed erythroid cells (Lammers *et al.*, 2002; Jayapal *et al.*, 2010). We confirmed that EPCAM (also known as CD326) was expressed in the majority of CD235a^+^cells at day 13 of iPSC differentiation (Figure 6B). We also detected a small number of CD326^+^CD235a^-^cells indicating that CD326 might be marking early erythroid commitment prior to CD235a acquisition (Figure 6B). To further explore this, we assessed the dynamics of these markers during the in vitro erythroid differentiation of umbilical cord blood CD34^+^(CB34^+^) cells. At day 10 of differentiation, CD326 is expressed in CD235^-^and CD235a^low^cells but not in CD235a^high^cells, that correspond to more mature erythroid cells (Figure 6B, Supplementary figure 4A). No CD326 expression was detected in cells at day 18 of the differentiation protocol (when the majority of cells are mature CD235a^+^cells) nor in the mature erythrocytes found in adult peripheral blood (Supplementary figure 4A). Taken together these data suggest that CD326 (EPCAM) marks early erythroid progenitors in both hiPSC-, foetal-and adult-derived cells.

**Figure 6.**
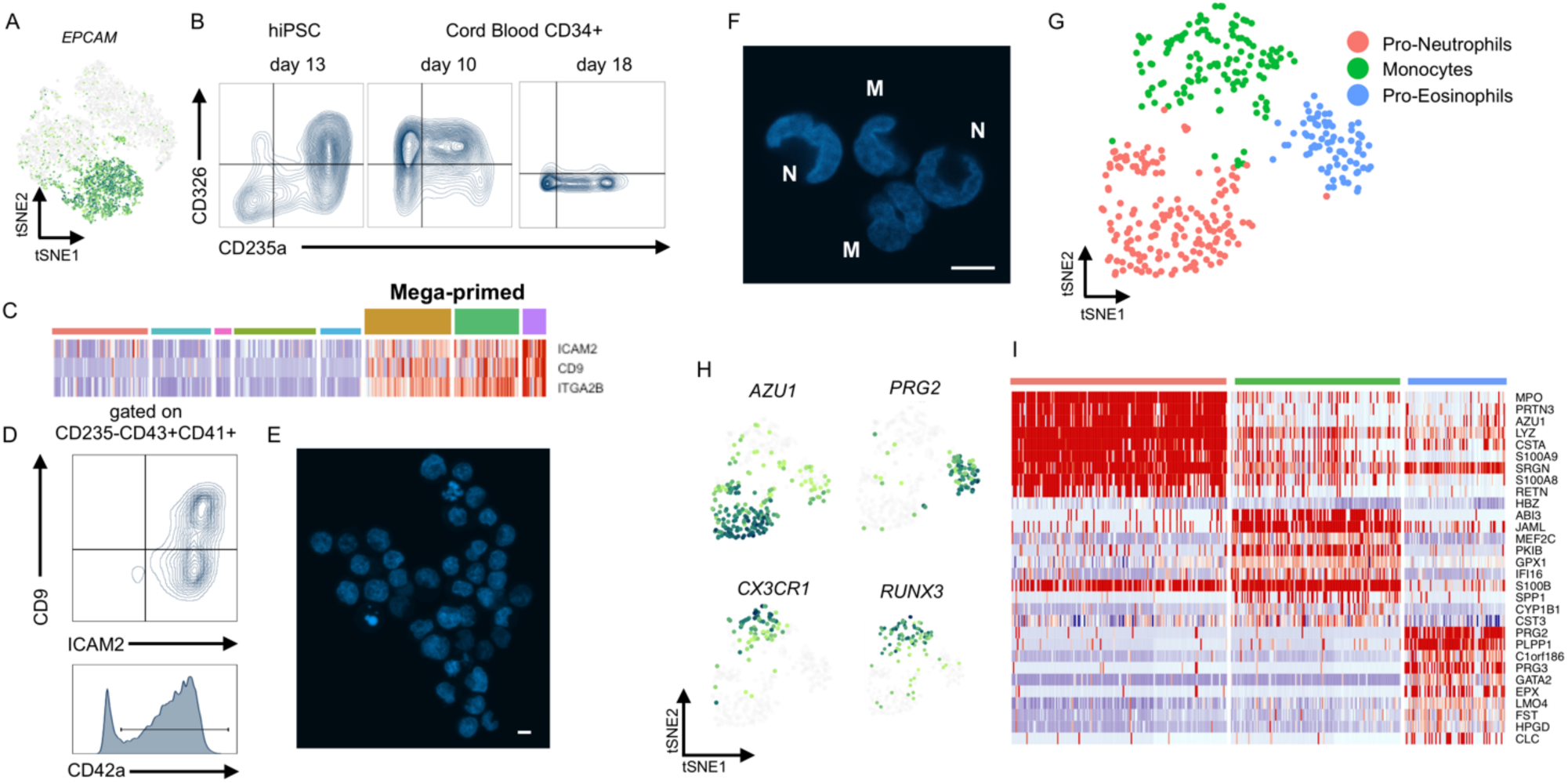
Primed clusters show immature lineage features of erythroid cells, megakaryocytes, neutrophils, eosinophils and monocytes. (**A**) *EPCAM* expression on tSNE. (**B**) Flow cytometry analysis of *EPCAM* expression hiPSC derived progenitors at day 13 and cord blood CD34+ differentiated in vitro. (**C**) Heatmap showing expression of ICAM2, CD9, and ITGA2B (also known as CD41). (**D**) Flow cytometry analysis of CD9 and ICAM2 expression in megakaryocytes biased progenitors gated on CD235a-CD43+CD41+, and CD42a expression in the same gated population. (**E**) DNA staining of sorted CD41+CD42+ cells and. (**F**) DNA staining of sorted CD235a-CD43+CD33+CD44-CD18+. (**G**) tSNE visualitation of subclustered ganulo-primed cells, showing 3 cell identities. (**H**) Gene expression of subclusters’ marker genes on tSNE. (**I**) Heatmap of the top 10 marker genes for each subcluster.

Three clusters with megakaryocyte and platelet signatures were identified (Mega-primed 1, 2 and 3), characterised by a high level of expression of *ITGA2B* (CD41), *GP9, PF4* (Figure 2C-D and Supplementary Table 1). We observed that the cell surface markers such as ICAM2 and CD9 seemed to be highly expressed later in their differentiation, in cluster Mega-primed 3 (Figure 6C). We confirmed the co-expression of these markers by flow cytometry and observed a population of CD41^+^CD9^+^ICAM2^+^cells, with around 85% of the CD41^+^cells also expressing CD42a (Figure 6D). When CD41^+^CD42^+^cells were sorted (Figure 6E), no polyploidy was detected supporting their immature stage (Figure 6F-Supplementary figure 4B).

In the mouse yolk sac, EMPs are distinguished from primitive haematopoietic progenitors by their capacity to initiate granulopoiesis, as indicated by morphological identification of neutrophils, eosinophils, basophils and mast cells (McGrath *et al.*, 2015). We identified a granulo-primed cluster in which granulocyte lineage markers such as *MPO, AZU1, RNASE2* together with *ITGB2* (coding for CD18) as a membrane marker (Figure 3C, Supplementary Table1). This observation, together with the potential of naïve 1 to give rise to granulocyte colonies, supports the definitive nature of the naïve 1 cluster, ascribable to an EMP-like progenitor. We analysed the nuclear morphology of sorted CD235a^-^CD43^+^CD18^+^CD33^+^CD44^-^cells by microscopy. As expected, we detected the existence of granulocytes and monocytes (and by inference, EMPs) by morphology (Figure 6G). To resolve this heterogeneity, we sub-clustered the data and identified three putative cell populations whose identity could be inferred by marker gene expression (Figure 6H-L). One of these sub-clusters co-expressed genes associated with the naïve unprimed progenitors (*LMO4, GATA2* and *FST*) as well as genes associated with the eosinophil lineage such as EPX, eosinophil peroxidase, and the proteoglycans PRG2 and PRG3 (Soragni *et al.*, 2015), and we speculate that these have a pro-eosinophil identity (Figure 6I-L). Another cluster appeared to express high levels of neutrophil-associated genes such as *MPO, AZU1, PRTN3, LYZ, S100A8* and *S100A9* and was thus annotated as a Pro-Neutrophil cluster (Figure 6I-L). Finally, we typed the third cluster as monocytes due to the expression of monocyte associated signature such as *CSF1R, CXC3R1*, and *IRF8* (Figure 6I-L). Noteworthy, *RUNX3* expression was specifically associated with the monocyte subcluster (Figure 6I). *RUNX3* role in developmental myelopoiesis has been shown in zebrafish, where treatment with RUNX3 morpholino lead to reduction of both stem cells and macrophages (Kalev-Zylinska *et al.*, 2003), and in mouse where it is expressed in Langerhans cells, a type of dendritic cell (Fainaru *et al.*, 2004). Tissue resident macrophages and dendritic cells are seeded by yolk sac monocytes that migrate to the embryo (Schulz *et al.*, 2012; Mass *et al.*, 2016; Stremmel *et al.*, 2018). Opposite to primitive macrophages, these cells develop from c-*myb*-expressing EMPs through a monocyte intermediate (Hoeffel *et al.*, 2015), further supporting our EMP-like identity of our human *MYB*^+^progenitors (supplementary table 1).

The lack of clusters with a fully differentiated transcriptome is in line with the fact that we did not employ terminal differentiation culture conditions and focused on the progenitor populations marked by CD43. These data show the competence of human iPSC-derived hematopoietic progenitors to initiate granulopoiesis in vitro and provide single cell transcriptome of human granulocyte progenitors.

## Discussion

We describe here the single cell transcriptome of human erythro-myeloid progenitors and their descendent primed cells. The priming direction observed from the naïve progenitors, confirmed by immunostaining and chimeric culture experiments, showed the ability of these cells to generate erythroid cells, megakaryocytes, monocytes, macrophages and granulocytes. This lineage output corresponds to that of mouse EMPs, definitive progenitors of the yolk sac (McGrath *et al.*, 2015; Frame *et al.*, 2016). Within the naïve progenitors, we observed high levels of *CD44* mRNA expression, a protein reported to be expressed in normal and leukemic hematopoietic cells (Zöller, 2015). Recently, a single cell transcriptome analysis reported that CD44 marks hematopoietic stem and progenitor cells in the mouse AGM, and functions in endothelium to hematopoietic transition, EHT (Oatley *et al.*, 2018). We confirmed CD44 expression on the aortic endothelium of the AGM region and, showed expression on some of the vessels in the yolk sac together with the hematopoietic clusters associated with these. From our functional assays, membrane expression of CD44 identifies human progenitors with clonogenic potential. Pseudotime and diffusion analyses showed that naïve 1, enriched for *CD44* expression, was at the top of the hierarchy, followed by naïve 2 and finally lineage primed clusters. To reveal the full lineage potential of progenitor cells we developed a chimeric culture system by co-culturing sorted ZeissGreen^+^population with untagged differentiating cells. When cells were exposed to the more complex microenvironmental cues, achieved in the differentiation, the lineage potential reflected by the priming signatures and the lineage output coincided. Despite the use of controlled differentiation conditions without feeder cell support or serum addition, the differentiating cells are a source of cytokines themselves, and this could explain the need of a system to trace lineage output of subpopulation while exposing the cells to the same stimuli from where the data set was obtained. Using chimeric tracing we also observed that progenitors are capable of moving between the naïve states, as well as progressing into primed states in a continuous fashion. Many studies employing single cell transcriptome and proteomic strategies appreciated a continuum of cell states as opposed to the sequential discrete cell types depicted in text-book hematopoietic hierarchies (Paul *et al.*, 2015; Velten *et al.*, 2017; Rodriguez-Fraticelli *et al.*, 2018; Knapp *et al.*, 2019). Our results showed that continuity is also associated with bidirectional fluidity between un-primed states suggesting a degree of cell plasticity, that during development could provide an advantage to face the changing demand of the growing embryo. The frequency of the naïve population was increased by BMP4 stimulation, and so pose a question on the role of BMP signalling on yolk sac EMP derived haematopoiesis, that has not been characterised so far. In the aorta-gonad-mesonephros, endothelial cells express BMP4 (Souilhol *et al.*, 2016), and display active BMP pathway with BMP-responsive element reporter (Crisan *et al.*, 2015). In contrast, within the HSC compartment BMP4 signalling needs to be inhibited (Souilhol *et al.*, 2016), via BMPER, for their full maturation (McGarvey *et al.*, 2017). Interestingly, here we showed that the naïve progenitors express high levels of Follistatin, a potent inhibitor of both BMP and TGFb, that could act as an autocrine/paracrine protection system against BMP-TGF signalling. Together with the observation of the BMP activity in the endothelial compartment, this may suggest that BMP4 could act by promoting the emergence of progenitors from the endothelium rather than expanding them. We show here that mouse and human progenitors co-express CD44 and LMO4, a LIM-domain protein widely expressed in the mouse embryo (Grutz, Forster and Rabbitts, 1998). LMOs form multiprotein complexes, as in the heptad complex where LMO2 binds other 6 transcription factors involved in HSPC (Wilson *et al.*, 2010), or in complexes involved in erythropoiesis (Wadman *et al.*, 1997). Recent sequencing experiments detected LMO4 expression in both adult mouse HSC (Lai *et al.*, 2018) and human granulocytes progenitors in the bone marrow (Paul *et al.*, 2015) but which proteins are bound to LMO4 in the progenitor compartment has yet to be identified. We also described high levels of ID2 and ID4 within the progenitors. IDs, like LMOs proteins, do not present DNA binding domain and rather act through binding of other proteins. This class of protein has not been extensively exploited in programming approaches as much as other transcription factors (Ivanovs *et al.*, 2017). Overexpression of IDs or LMOs could not only provide an alternative approach for programming gene cocktails, but it could also be employed to maintain the progenitor state in culture and prevent their differentiation. In support of this idea, overexpression of ID2 in human HSC from cord blood has been reported to enhances their functional stemness in vivo (van Galen *et al.*, 2014).

Taken together our data show that cytokine stimulation of hPSCs in vitro recapitulate extraembryonic haematopoiesis with erythro-myeloid progenitors and describe CD44 as a marker for human EMPs. We identified the transcriptome of naïve and primed human progenitors in vitro, which represent a potential source for both monocytes/macrophages and granulocytes for therapeutic application and provides an important resource for the identification of target genes for HSC generation in vitro.

## Acknowledgment

The work was funded by Wellcome Trust (Grant No. 102610), MRC Innovate UK (Grant No. 102853), BBSRC (Grant No. S002219/1). AF received a Carnegie Incentive Grant (Grant No. RIG008218). Sequencing and following alignment was carried out by Edinburgh Genomics, The University of Edinburgh. Edinburgh Genomics is partly supported through core grants from NERC (R8/H10/56), MRC (MR/K001744/1) and BBSRC (BB/J004243/1). We thank Andrew Elefanty for sharing the RUNX1C-GFP cell line. We thank Fiona Rossi, Claire Cryer, Bindi Heer from the Flow Facility, Bertand Verney and Matthieu Vermeren from the imaging facility.

## Author Contribution

FA, designed and performed research, analyzed the data and wrote the manuscript. NR performed bioinformatics analysis. PR, ST, MLY, AHT, JE, BH, RA performed research. LMF designed the experiment, analyzed data and wrote the manuscript. NH, AM, KO, provided intellectual input and final approval of the manuscript.

## Methods

### Pluripotent stem cells culture

hPSCs were maintained in vitro in StemPro hESC SFM (Gibco) with bFGF (R&D) at 20 ng/ml for SFCi55 and KLF1-mCherry-SFCi55, and at 40 ng/ml for H9 and RUNX1C-GFP. Wells were coated with CELLstart at least 1 hour before plating and cells were passaged using the StemPro EZPassage tool (ThermoFisher Scientific). Media change was performed every day and cells passaged every 3–4 days at a ratio of 1:4.

### hPSCs hematopoietic differentiation

hPSCs were differentiated in a xeno-free composition of SFD medium (Sturgeon *et al.*, 2014), BSA was substituted with human serum albumin, HSA, (Irvine-Scientific). Day 0 differentiation medium, containing 10 ng/ml BMP4 was added to the colonies prior cutting. Cut colonies were transferred to a bacteriological grade well to form embryoid bodies and cultured for two days. At day 2 media was changed and supplemented with 3 µM CHIR (StemMacs). At day 3, embryoid bodies were collected and dissociated in Accutase (Gibco) to single cell solution, cells were plated on tissue culture grade wells in SFD medium supplemented with 5 ng/ml bFGF and 15 ng/ml VEGF (2×10^5^cells/well on 6 well plates or 1×10^5^cells/well on 12 well plate). At day 6 media was changed for final hematopoietic induction in SFD medium supplemented with 5 ng/ml bFGF, 15 ng/ml VEGF, 30 ng/ml IL3, 10 ng/ml IL6, 5 ng/ml IL11, 50 ng/ml SCF, 2 U/ml EPO, 30 ng/ml TPO, 10 ng/ml FLT3L and 25 ng/ml IGF1. From day 6 onward, cytokines were replaced every two days. For further erythroid differentiation, SFD2 was used (IMDM, 10% HAS, 10 ng/ml insulin (Sigma-Aldrich), 200µ/ml Human Holo-Transferrin and Glutamax). At day 13, 3 × 10^5^cells were replated in SFD2 supplemented with 50 ng/ml SCF, 16.7 ng/ml FLT3L, 6.7 ng/ml BMP4, 6.7 ng/ml IL3, 6.7 ng/ml IL11, 3U/ml EPO, 50 µM IBMX and 10 µM Hydrocortisone. At day 20, 10^6^cells were replated in SFD2 supplemented with 3U/ml EPO, 6.7 ng/ml IL3, 6.7 ng/ml IL11, 20 ng/ml SCF and IGF1 20 ng/ml. From day 27 to day 30, 10^6^cells were terminally differentiation in SFD2 medium with 3U/ml EPO.

### Pluripotent stem cell targeting

Regulatory region spanning the transcriptional starting site of KLF1 gene was amplified using PrimeStarMAX (Takara) with the addition of KpnI sites. This 1.4 KB region, spans the distal promoter (−790) to the intronic enhancer (+600) of the KLF1 gene contain (Siatecka *et al.*, 2010). MCherry tag with polyadelintion signal was amplified using the same strategy and with the addition of flanking KpnI and EcoRI sites. PCR product were assembled into KpnI and EcoRI difgested pZDonor-AAVS1 Puromycin plasmid (Sigma-Aldrich) and ligated using the Quick Ligase Kit (New England Biolab). Correct clones were identified by sequencing. For targeting of hiPSC line SFCi55, 107 cells were electroporated with 40µg of targeting vector and 5 µg of AAVS1-ZNF-Left and µg 5 AAVS1-ZNF-Right. Cells were grown under Puromycin selection at 0.2 µg/ml for at least 4 weeks. Colonies were manually picked and expanded, genomic DNA was purified and genotyped according to manufacturer instruction in the pZDonor-AAVS1 Puromycin Kit. Correctly integrated clones were tested in differentiation condition and the clone #9 was selected for further analysis.

### Cord blood erythroid differentiation

Frozen Umbilical cord blood (UCB) derived CD34^+^cells were purchased from Stemcell Technologies (Cat No. 70008.5) from consenting donors with protocols approval by either the Food and Drug Administration (FDA) or an Institutional Review Board (IRB). Cells were plated at 1-6 × 10^4^cells/ml in ISHIT base media (Iscove’s Basal Media (Biochrom AG), 5% human AB^+^serum, 3 U/ml heparin and 10 mg/ml Insulin) supplemented with 60 ng/ml SCF, 5 ng/ml IL3, 3 U/ml EPO, 1mM Hydrocortisone and 200 mg/ml human holo-Transferrin. At day 6, batches were frozen for further use at 10^6^cells/ml in 60% ISHIT media, 30% knockout serum replacement (Gibco) and 10% DMSO. Cells where thawed and cultured for additional 2 days in the same medium. At day 8, cell density was adjusted to 10^5^cells /ml in ISHIT media supplemented with 10 ng/ml SCF, 3 U/ml EPO, 300 mg/ml and human holo-Transferrin, and cultured for a further 3 days. Finally, cells were cultured at a density of 10^6^cells/ml in ISHIT medium supplemented with 3U/ml EPO and 300 mg/ml human holo-transferrin until day 21. Media was changed every 3-4 days throughout the protocol.

### Flow cytometry staining and cell sorting

From hematopoietic differentiation, suspensions cells were collected from the well by aspiration of the media, adherent cells were detached from the well by using Cell Dissociation Buffer (ThermoFisher). Cells were washed with PBS + 1% BSA, counted and were stained at 10^5^cells for a single tube. Cells were stained with antibodies in supplementary table X, for 30’ at RT gently shaking. Flow cytometry data were collected using DIVA software (BD). Sorting was performed using FACSAria Fusion (BD) and cells were collected in PBS + 1%BSA. Data were analysed using FlowJo version 10.4.2.

Yolk sacs and AGM from mouse embryos were micro dissected in PBS supplemented 2% FBS and washed twice before tissue digest. Single cell suspensions were obtained by incubation in 0.125% collagenase at 37°C for 45’ for AGMs and 60’ for yolk sacs, followed by mechanical dissociation by pipetting and a final wash.

### Antibody panels

The following antibodies were used for multicolour panels. Figure 1 and figure 2: CD235a-PeCy7 (1:100, BD, GA-R2(HRI2)), CD43-APC (1:100, eBiosceince, eBioB4-3C1) and DAPI. Figure 4 A, E: CD235a-FITC (1:100, eBioscience, HRI2(GA-R2)), CD43-APC (1:100, eBiosceince, eBioB4-3C1), CD33-Pecy7 (1:200, BioLegend, WM53), CD44-PB (1:50, BioLegend, BJ18), CD18-Pe (1:100, BioLegend, 1B4/CD18) and DAPI. Figure 4 D: CD235a-FITC (1:100, eBioscience, HRI2(GA-R2)), CD43-APC (1:100, eBiosceince, eBioB4-3C1), CD33-Pecy7 (1:200, BioLegend, WM53), CD44-PB (1:50, BioLegend, BJ18), and DAPI. Figure 4 B,C, and F: CD235a-BV605 (1:300, BD, GA-R2(HRI2)), CD43-APC (1:100, eBiosceince, eBioB4-3C1), CD33-Pecy7 (1:200, BioLegend, WM53), CD44-PB (1:50, BioLegend, BJ18), CD18-Pe (1:100, BioLegend, 1B4/CD18) and DAPI. Figure 5: CD44-PE (eBioscience; IM7, 1:200), CD45-BV421 (Biolegend; 30-F11, 1:200) and Draq7. Figure 6 A: CD326-BV785 (1:100, BioLegend, 9C4), CD235a-FITC (1:100,eBioscience, HRI2(GA-R2)) and DAPI. Figure 6F Figure 4 D: CD235a-FITC (1:100, eBioscience, HRI2(GA-R2)), CD43-APC (1:100, eBiosceince, eBioB4-3C1), CD33-Pecy7 (1:200, BioLegend, WM53), CD44-PB (1:50, BioLegend, BJ18), and DAPI. Figure 6D: CD235a-FITC (1:100, eBioscience, HRI2(GA-R2)), CD43-APC (1:100, eBiosceince, eBioB4-3C1), ICAM2-Pe (1:100, BioLegend, CBR-IC2/2), CD9-APC-Fire750 (1:100, BioLegend, HI9A), CD41-PB (1:50, BioLegend, HIP8) and Darq7 (ThermoScietific), and CD41-PB (1:50, BioLegend, HIP8) and CD42-Pe (1:100, BD, ALMA 16).

### Methylcellulose assay

Sorted cell populations were counted and plated at 5000 cells into 2 ml of methylcellulose medium (Human enriched H4435, Stemcell Technologies). Cells were incubated in the assay for 14 days and then scored.

### Single cell RNA sequencing

Two independent sample of hiPSC SFCi55 were differentiated synchronously and sorted at day 13 using CD235a^-^CD43^+^immunophenotype, viable cells were sorted using DAPI. Cell viability was also confirmed by Trypan blue stain for accurate count using TC20 cell counter (Biorad). Around 12000 cells per sample were loaded into the 10X Chromium Controller and single cell libraries were obtained using the Chromium single cell 3’ Reagent Kits v2 (10XGenomics) according to manufacturer protocol. RNA concentration was obtained using Quibit RNA HS (Thermo-Fisher). Quality of the obtained libraries were verified using LabChip GX (PerkinElmer). Libraries were sequenced using HiSeq 4000 technology (Illumina) at 50000 reads/cell. Data were aligned to GRCh38 using the Cell Ranger dedicated pipeline (10XGenomics). Data filtering, dimension reduction, clustering analysis and differentially expressed genes were obtained using Seurat.R package, cell trajectories was obtained using Monocle.R (code and data are available on request through corresponding authors).

### Mouse embryo and yolk sac embedding and sectioning

Whole embryos with the yolk sac were fixed in 4% PFA overnight at 4°C on a gentle shaker, yolk sac was detached from the embryo after fixation and processed in parallel to the embryo. After rinsing them twice with PBS, they were placed in a solution of 15% sucrose/PBS for 3h at 4°C and then transferred into PBS with 15% sucrose and 7% gelatine at 37°C for 1-3h until they sank. Embryos and yolk sacs were mounted in gelatin using mounting specimens (Sigma-Aldrich), snap-frozen in liquid nitrogen and stored at - 80°C. 7μm-thick sections were cut using Cryotome FSE (Thermo ScientifiC) and further processed for immunostaining or stored at -20°C

### Immunostaining and microscopy

Gelatin was removed boiling the slide for 30” in PBS and washed twice in PBS, permeabilized and blocked with 5% donkey serum + 0.3% triton-X100/PBS and stained with primary antibodies overnight at 4°C (CD44: Stratech KM201, 1:100; LMO4: Thermo-Fisher PA5-24248, 1:200). Following three washes in PBS, sections were stainied with secondary antibodies in 0.3% triton-X100/PBS for 2 hours at room temperature (anti-rat Alexa Fluor 568, Thermo Fisher A-11077, 1:500, anti-rabbit Alexa Fluor 488, Thermo Fisher A-11008, 1:500). After three washes in PBS, slides were counterstaining in 10 mg/ml DAPI. Samples were mounted with ProLong Gold Antifade mountant (Life Technologies) and dried at room temperature in the dark for a minimum of 2h, then stored at 4°C. Images were acquired using an inverted confocal microscope (Leica SP8) and analyzed using ImageJ 1.50i (https://imagej.nih.gov/ij).

### Statistical analysis

All data are reported as mean ± standard error mean (SEM), statistical tests and p values are reported within figure legends. Statistical analysis was performed with Graph Pad Prism version 6 and R.

