## Supplementary figures for "Single cell transcriptome analysis reveals markers of naïve and lineage-primed hematopoietic progenitors derived from human pluripotent stem cells"

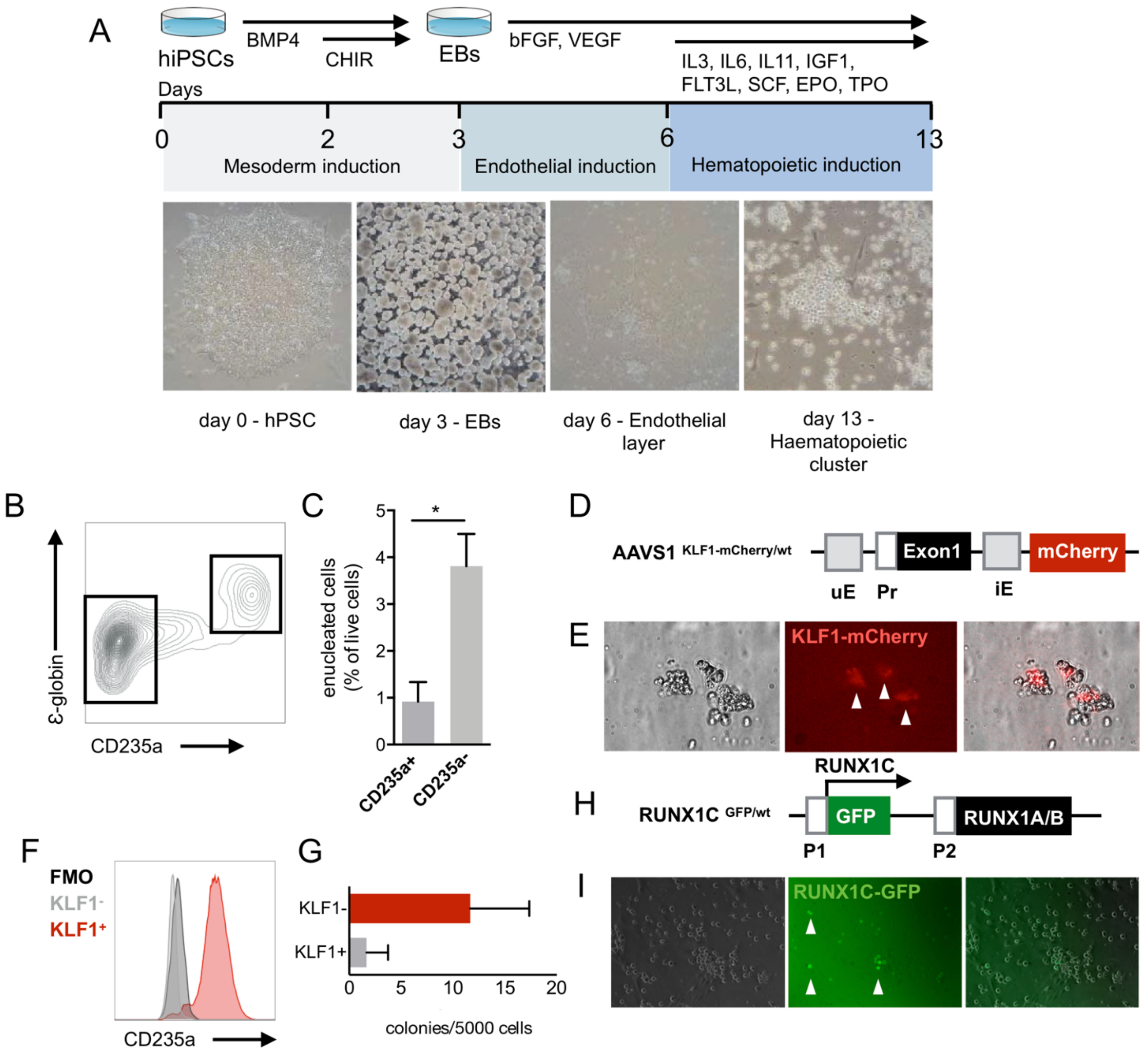


**Supplementary Figure 1 - Human definitive hematopoietic progenitors resides in the CD235a- compartment.**

(**A**) Schematic of the differentiation protocol. (**B**) Representative expression profile of embryonic globin (ε-globin) and CD235a. (**C**) Enucleation efficiency of sorted CD235a+ and CD235a- cells in erythroid differentiation conditions (n=3, $p<0.05$, paired t-test). (**D**) Representative AAVS1 locus structure of KLF1-mCherry human iPSC reporter cell line. 1.4 Kb region spanning the TSS of *KLF1* containing the upstream enhancer (uE), proximal promoter (Pr), and intronic enhancer iE upstream the mCherry tag. (**E**) primitive erythroid colonies expressing KLF1 in methyl cellulose assay are pointed by arrowheads. (**F**) Distribution of KLF1+ and KLF1- in relationship to CD235a. (**G**) Colony forming potential of sorted KLF1+ and KLF1- cells (n=3, ns, paired t-test). (**H**) Representative locus structure of RUNX1C-EGFP human ESC cell line (P1 - distal promoter, P2 - proximal promoter). (**I**) Differentiated cells expressing RUNX1C are pointed by arrowheads.

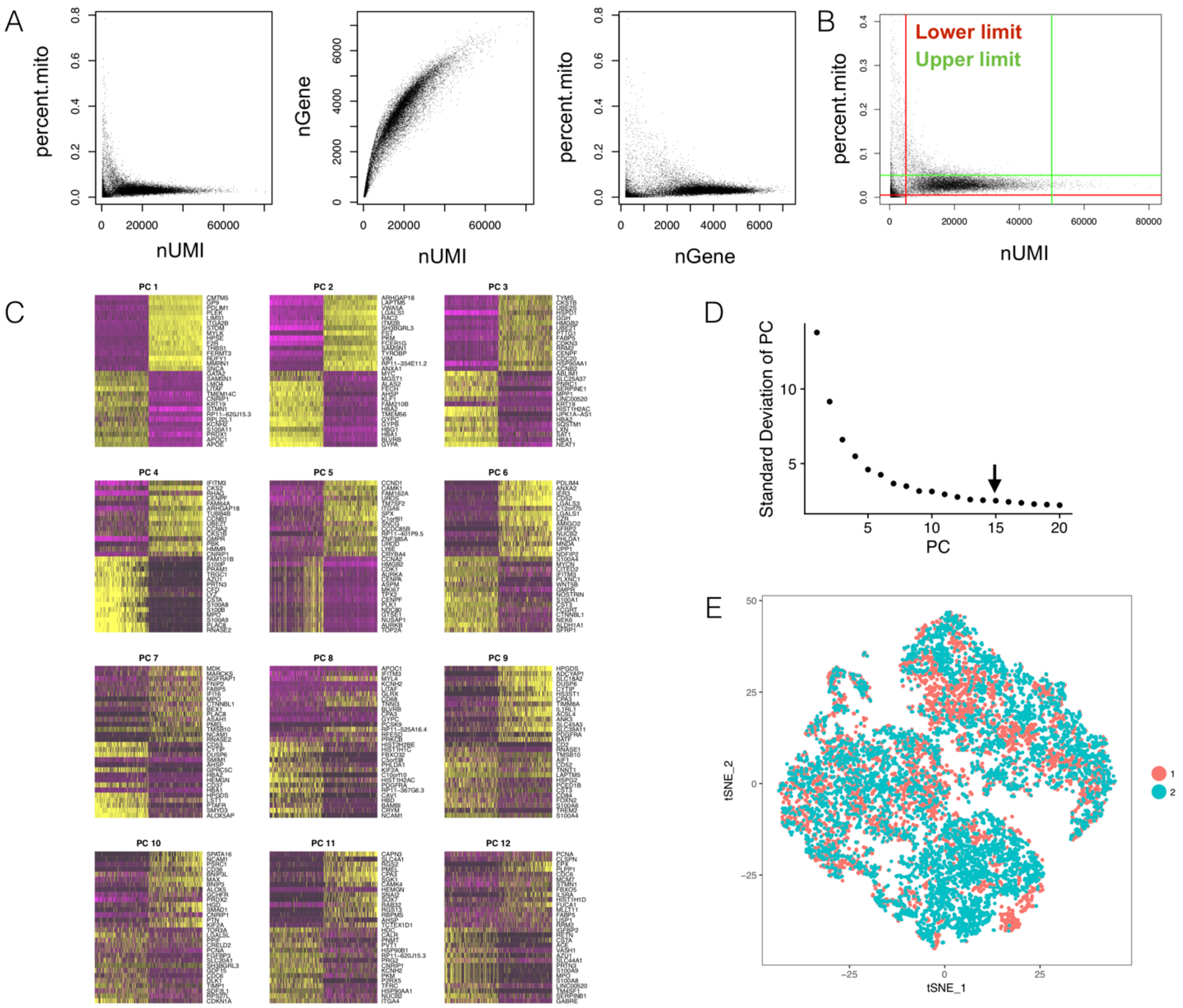


**Supplementary figure 2 – Quality controls of the single cell RNA sequencing experiement.**

(**A**) Scatter plot showing data for each unfiltered cells of mitochondrial genes (percent.mito), number of detected genes (nGene) and the numer of unique molecule identifier (nUMI). (**B**) Limit used to filter cells based on mitochondrial genes and UMI counts. (**C**) Heatmap of genes in the first 12 principal component. (**D**) Elbow plot explaining the standard deviation described by each principal component (PC), arrow indicates the number of PC used for further analyses. (**E**) tSNE plot of two different sample 1 and 2 showing overlapping cells.

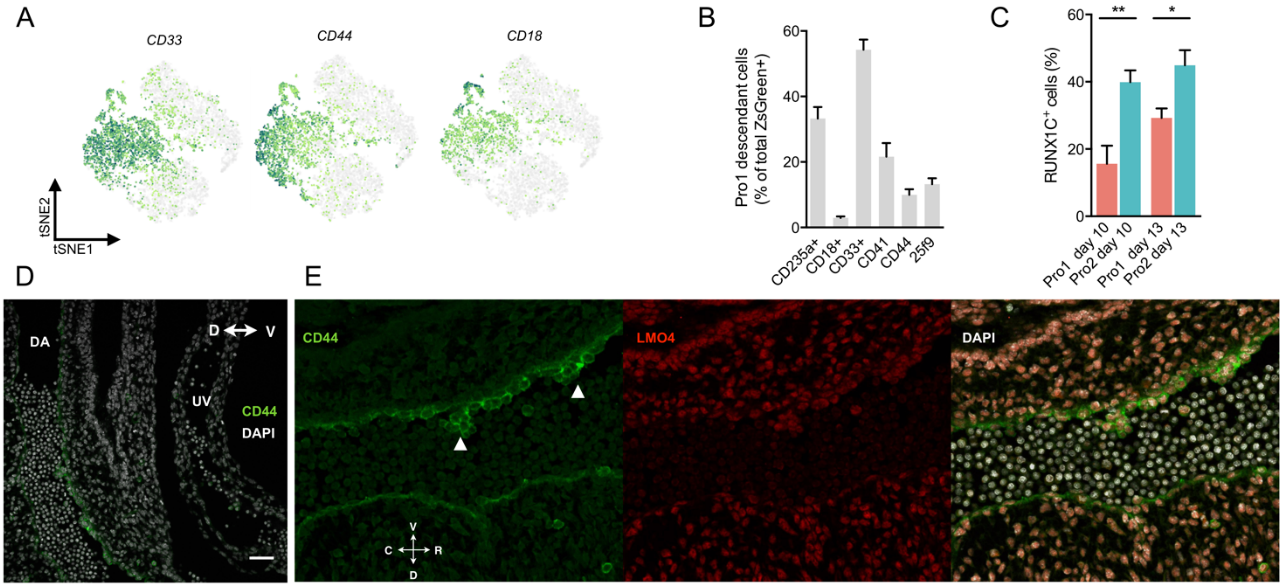


**Supplementary figure 3 – Functional validation of progenitors clusters.**

(**A**) Gene expression of *CD33*, *CD44* and *CD18* on tSNE. (**B**) Naive 2 descendant cells into different lineages, quantification of flow cytometry single stains (n=4-7). (**C**) RUNX1-GFP expression in Naive 1 and Naive 2 at both day 10 and day 13 (n=12, paired t test; * p<0.05, ** p<0.01) (**D-E**) CD44 and LMO4 immunostaining in mouse sections of the AGM at day E10.5 (DA - dorsal aorta, UV - umbilical vein, arrowhead points at intra-aortic hematopoietic cluster).

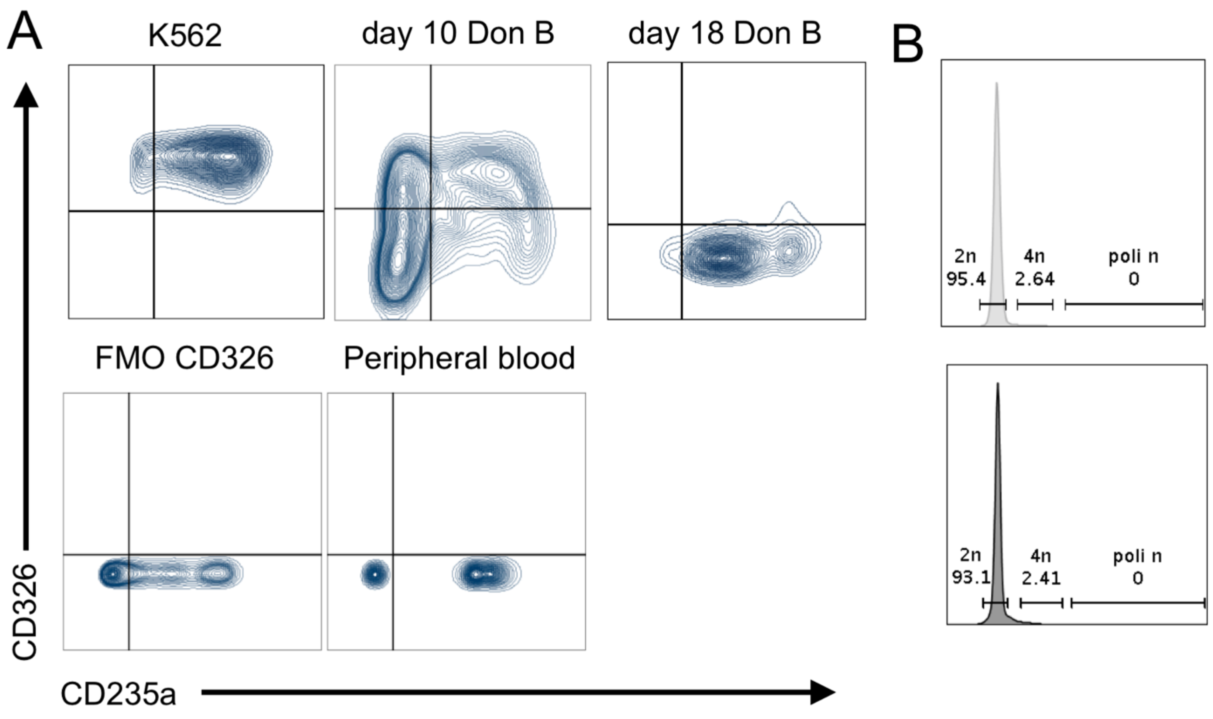


**Supplementary figure 4 – Primed progenitors show immature features.**

(**A**) Membrane expression of CD326 in relationship with CD235a in positive control cells (K562), Cord blood derived erythroid cells (day 10 and day 18), in peripheral blood and in negative control (FMO). (**B**) Flow cytometry analysis of DNA content in sorted CD41$^{+}$CD42$^{+}$(light grey) compared to unsorted cells (dark grey).

**Supplementary Table 1 – Clusters'markers.**

List of markers' genes for each cluster ordered by average log fold change (avg_log_FC) with respective adjusted p value s (adj_p_val).
